# DISRUPTION OF A DNA REPAIR PROTEIN PROMOTES ANTIBIOTIC RESISTANCE IN *ACINETOBACTER BAUMANNII*

**DOI:** 10.64898/2026.08.27.747542

**Authors:** Suman Tiwari, Syed H. Raza, Namrata Bonde, Roberto Jhonatan Olea-Ozuna, Tuhina Maity, Muneer Yaqub, Tahira Amdid Ratna, Kelli Palmer, Joseph Boll, Jonathan Monk, Nicholas A. Dillon

## Abstract

*Acinetobacter baumannii* is a high-priority Gram negative opportunistic pathogen known for its high rates of multidrug resistance (MDR). Minocycline (MIN), a tetracycline class antibiotic, is one of the most effective antibiotics for treating *A. baumannii* infections in patients. Unfortunately, MIN resistance is spreading internationally and has begun to emerge in the United States. While efflux pumps are correlated with MIN resistant *A. baumannii*, clinical data suggests alternative mechanisms of MIN resistance. To explore the genetic basis for MIN resistance in *A. baumannii* we employed a machine learning model to predict genetic resistance correlates from clinical isolates. Mutations in *ruvB*, a DNA repair protein, were strongly correlated with MIN resistant clinical strains of *A. baumannii*. Consistent with the prediction, tn26 insertion in *<u>ruvB</u>* in *A. baumannii* strain AB5075, and deletion of *ruvB* in strain ATCC 19606, increased MIN minimum inhibitory concentrations to a level that exceeds the MIN resistance breakpoint. RuvB complexes with RuvA and RuvC to resolve Holliday junctions during recombination. However, only *ruvB* mutants showed the resistance phenotype; neither *ruvA* nor *ruvC* mutants were MIN resistant, suggesting loss of the complex’s activity was not the basis for resistance. We observed *ruvB* mutants produced increased biomass during planktonic growth relative to the other two *ruv* mutants. Upon examination, the *ruvB*::tn26 mutant had a 451% increase in biomass and 360% thicker biofilms relative to wildtype. We determined the disruption of *ruvB* lead to thicker biofilms and enriched in extracellular DNA (eDNA), and DNase I treatment collapsed the enhanced biofilm phenotype and markedly reduced tetracycline-class MICs. FLAG–RuvA accumulated within the biofilm matrix in the absence of RuvB, supporting a model in which RuvA contributes to stabilization of eDNA-rich structures. In a murine pneumonia model, *ruvB* disruption did not significantly alter survival or pulmonary burden in untreated infection but reduced bacterial dissemination and increased minocycline resistance. Together, these findings reveal an unexpected connection between Holliday junction processing, eDNA-rich biofilm architecture, and antibiotic resistance in *A. baumannii*.

## Introduction

Antimicrobial resistance (AMR) is a mounting global health crisis, responsible for an estimated 4.95 million deaths in 2019, with Gram-negative organisms disproportionately contributing to this burden(1). Among them, *Acinetobacter baumannii* has emerged as one of the most formidable members of the ESKAPE pathogens. This opportunistic bacterium is a frequent cause of ventilator-associated pneumonia, bloodstream infections, and wound infections in critically ill patients, where mortality rates often exceed 40% in multidrug-resistant (MDR) cases(2, 3). Its ability to withstand desiccation, tolerate disinfectants, and persist on hospital surfaces enables transmission in healthcare environments and contributes to its recognition as a global priority pathogen.

Therapeutic options for MDR *A. baumannii* are limited. Minocycline, a semi-synthetic tetracycline, has retained notable efficacy, with susceptibility rates exceeding 70% in surveillance studies(4, 5). However, resistance to tetracyclines continues to emerge, primarily driven by efflux pumps, ribosomal protection proteins, and, increasingly, adaptive phenotypes such as biofilm formation(6). Biofilms, complex multicellular aggregates encased in extracellular matrices of polysaccharides, proteins, and extracellular DNA (eDNA), provide both physical and physiological protection against antimicrobial agents. Within biofilms, bacteria often exhibit antibiotic resistance that exceeds planktonic cells, and robust biofilm formation in *A. baumannii* clinical isolates has been strongly associated with multidrug resistance and treatment failure(7).

To move beyond known mechanisms and uncover novel determinants of resistance, we developed a machine-learning framework trained on genomic and phenotypic data from over 1,400 *A. baumannii* clinical isolates. This model successfully recapitulated established pathways, such as tetracycline efflux and ribosomal protection, thereby validating its predictive capacity. Strikingly, it also identified the gene *ruvB*, which encodes the ATPase subunit of the RuvABC Holliday junction branch migration complex, as one of the strongest predictors of resistance. In addition to minocycline resistance (p = 0.0061), *ruvB* variants were significantly associated with reduced susceptibility to doxycycline and vancomycin, and correlated with resistance across multiple drug classes including amikacin (p = 0.015), ampicillin-sulbactam (p = 0.0003), ciprofloxacin (p = 0.039), doripenem (p = 2.76 × 10⁻⁸), and imipenem (p = 1.81 × 10⁻⁵). These findings were unexpected given the canonical role of RuvB in DNA repair. The RuvABC complex ensures the resolution of Holliday junctions during homologous recombination, safeguarding genome integrity under stress. Yet our computational analysis suggested that the function of *ruvB* may extend beyond DNA metabolism, raising the possibility that this repair pathway influences antibiotic resistance through mechanisms not previously appreciated in *A. baumannii*.

The RuvABC complex is a mediator of homologous recombination, functioning at the core of bacterial DNA repair mechanism. RuvA recognizes and binds Holliday junction intermediates, RuvB provides ATP-dependent branch migration, and RuvC cleaves the junction to restore linear duplex DNA. This pathway is critical for resolving stalled replication forks and repairing DNA damage, thereby ensuring genome stability under conditions of stress. Historically, DNA repair systems such as RecA-mediated recombination and mismatch repair have been studied for their influence on mutation rates and evolutionary adaptation, with defective repair accelerating the emergence of resistance in organisms like *Escherichia coli* and *Pseudomonas aeruginosa*(8). DNA repair proteins have also been implicated in stress-induced tolerance, where their activation mitigates the bactericidal effects of fluoroquinolones and other DNA-damaging agents(9, 10). Yet despite these links, a direct role for repair proteins in shaping biofilm physiology or antibiotic resistance has not been defined in *A. baumannii*.

Intriguingly, recent work has revealed a potential conceptual bridge. In *E. coli* and other species, RuvA has been shown to substitute for DNABII-family proteins in stabilizing extracellular DNA lattices within biofilms, where branched DNA structures serve as architectural scaffolds(11). This finding suggests that components of recombination machinery, classically understood as guardians of genome integrity, may adopt unanticipated structural roles in the extracellular matrix. In this light, disruption of *ruvB* could free RuvA from its canonical partner, enabling it to stabilize eDNA within the biofilm matrix. Such a shift would provide a mechanistic basis for the enhanced biofilm formation and broad-spectrum resistance predicted by our computational analysis. However, whether this paradigm applies to *A. baumannii* has not been investigated, leaving a critical gap in our understanding of how DNA repair intersects with antibiotic resistance in this high-priority pathogen.

Here, we set out to test the hypothesis that disruption of *ruvB* promotes broad-spectrum antibiotic resistance in A. baumannii by stabilizing eDNA and enhancing biofilm formation. Although ruvB emerged as the strongest candidate from our computational analysis, we extended our investigation to the broader RuvABC system. In AB5075, we analyzed ruvA, ruvB, and ruvC transposon insertion mutants from the Manoil library and performed genetic complementation. In parallel, we constructed clean single (ruvA, ruvB, ruvC) and double (ruvAB, ruvAC, ruvBC) deletion mutants in ATCC 19606 using recombineering, alongside complementation in both strain backgrounds. This systematic approach allowed us to define the contribution of the entire RuvABC complex to antibiotic resistance and biofilm physiology. Using these complementary genetic and phenotypic tools, we reveal a previously unrecognized role for DNA repair proteins in biofilm-mediated resistance and establish ruvB disruption as a paradigm of broad antibiotic resistance in *A. baumannii*.

## Methods

### Bacterial Strains and Growth Conditions

*Acinetobacter baumannii* AB5075-UW (wild type and ruvA::tn26, ruvB::tn26, ruvC::tn26 mutants) was obtained from the Manoil transposon library (University of Washington). AB5075 is recognized as a model organism for MDR *A. baumannii* due to its virulence in multiple infection models and availability of curated mutant resources(12, 13). We further validated AB5075 as a model strain by comparing its MIC profile against a panel of clinical isolates (see Results). In parallel, ATCC 19606 was used to generate clean, markerless single and double deletions (ΔruvA, ΔruvB, ΔruvC, ΔruvAB, ΔruvAC, ΔruvBC) using a RecAB recombineering system adapted for A. baumannii (14, 15).

Unless otherwise stated, genetic manipulations (mutagenesis, cloning, electroporation) were performed in LB at 37 °C, whereas phenotypic assays (MIC testing, biofilm quantification, confocal microscopy) were conducted in cation-adjusted Mueller-Hinton broth (CA-MHB) following CLSI M100 guidelines(16, 17). Antibiotic concentrations were kanamycin 25 µg/mL, tetracycline 10 µg/mL, hygromycin 250 µg/mL (AB5075) or 400 µg/mL (19606) for genetic manipulations.

### Mutagenesis and Markerless Deletion in ATCC 19606

Markerless single (ΔruvA, ΔruvB, ΔruvC) and double mutants (ΔruvAB, ΔruvAC, ΔruvBC) were generated using a RecET-based recombineering system adapted for *A. baumannii* (14). ATCC 19606 carrying pMMB67EH-RecAb (Tet 10 µg/mL) was induced with 2 mM IPTG, harvested at OD₆₀₀ ≈ 0.5, and washed in ice-cold 10% glycerol. PCR donor fragments containing FRT-flanked kanamycin cassettes were amplified from pKD4 using primers with ∼125 bp flanking homology. Cells were electroporated (2-mm cuvette, 1.8 kV), recovered for 1 h in LB + IPTG and plated on LB agar + 25 µg/mL kanamycin. Recombinants were validated by colony PCR with primers external to the recombination junction. For marker excision, kanamycin-resistant colonies were transformed with pMMB67EH-FLP (Tet 10 µg/mL) and induced with IPTG to activate FLP recombinase. Tetracycline markers were subsequently removed by counterselection on LB agar + 2 mM NiCl₂ for 72 h. Colonies were streak-purified on LB agar without antibiotics to ensure plasmid curing. Final clean, markerless deletions were confirmed by PCR(18, 19).

### Complementation Strategy

For complementation, ruvA, ruvB, and ruvC coding regions were amplified with primers containing engineered BamHI (5′) and SalI (3′) sites, digested, and ligated into pJMP3665HygR. Constructs were verified in E. coli DH5α, then electroporated into AB5075 or ATCC 19606. Transformants were selected on hygromycin (250 µg/mL for AB5075; 400 µg/mL for ATCC 19606) and confirmed by colony PCR. Complementation restored wild-type phenotypes in MIC and biofilm assays. Primers are listed in Table S1.

### Minimum Inhibitory Concentration (MIC) Testing

MICs were determined by broth microdilution in CA-MHB using a high-throughput OT-2 robotic pipeline with automated Python endpoint analysis, validated against manual CLSI microdilution with 100% concordance(20). Overnight cultures were diluted to ∼5 × 10⁵ CFU/mL, dispensed into 96 well plates, incubated at 37 °C for 18-20 h, and OD₆₀₀ measured to assign endpoints.

### Biofilm Formation Assays

Static biofilms were quantified using the crystal violet microtiter plate assay(21). Cultures were diluted 1:100 in CA-MHB, incubated in 96-well PVC plates. After 24 h static incubation at 37 °C, wells were washed three times with PBS, stained with 0.1% crystal violet for 15 min, rinsed, and air-dried. Bound stain was resolubilized with 30% acetic acid, and absorbance was measured at 550 nm using a BioTek Synergy H1 plate reader. Each condition included eight technical replicates and ≥3 biological replicates.

### Confocal Microscopy of Biofilms

Biofilms were grown on ethanol-sterilized glass coverslips in CA-MHB for 72 h with media changing after every 24hrs, fixed in 4% paraformaldehyde (45 min), and sequentially stained with BODIPY FL NHS Ester (0.125 µg/mL in PBS, 5 min) to label neutral lipids (22, 23), quenched with 50 mM NH₄Cl (5 min); Texas Red hydrazide (2 µg/mL, 15 min) for proteins(24), Hoechst 33258 (2 µg/mL, 10 min) for DNA(11), and Sytox Red (2 µg/mL, 15 min) for eDNA(25, 26). Slides were mounted in ProLong™ Diamond and imaged on a Zeiss LSM 880 using sequential excitation (405/488/561/682 nm).

### Construction and visualization of FLAG-tagged RuvA in biofilms

To visualize RuvA localization within biofilms, a ruvAB double mutant was constructed in *A. baumannii* ATCC 19606 background as described above. For complementation of double mutant, FLAG-tagged ruvA was cloned into the BamHI/SalI sites of pMMB67EHKanR (kan 25 µg/mL). For control strains, the ruvAB mutant was further complemented with an untagged ruvB (hygromycin, 400 µg/mL) to restore the intact RuvAB complex as described above.

Biofilms were grown on glass coverslips and fixed prior to staining. DNA was visualized using Hoechst dye, while FLAG-tagged RuvA was detected using a recombinant rabbit monoclonal anti-FLAG (DYKDDDDK) antibody conjugated to Alexa Fluor™ Plus 594. Confocal Z-stacks were acquired under identical imaging conditions for all strains. Wild-type strains lacking FLAG-tagged constructs served as negative controls to assess background fluorescence.

### Murine pneumonia model

All animal experiments were approved by the Institutional Animal Care and Use Committee (IACUC) at the University of Texas at Dallas and conducted in accordance with institutional and federal guidelines. Female C57BL/6 mice (6-8 weeks old) were anesthetized with ketamine/xylazine (10:1) and intratracheally inoculated with A. baumannii AB5075 wild-type or ruvB mutant strains. Inocula were prepared from mid-logarithmic phase cultures, washed, and resuspended in sterile phosphate-buffered saline (PBS). Mice received a 1 x 10^8^ CFU bacterial dose in a total volume of 40 µL(27). For survival studies, animals were monitored at regular intervals for morbidity and mortality for up to six days post-infection. For bacterial burden analysis, mice were euthanized at 30 hr time point post-infection, and lungs were aseptically harvested, homogenized, serially diluted, and plated for enumeration of colony-forming units (CFU)(27).

For antibiotic treatment studies, C57BL/6 mice were intratracheally infected with AB5075 wild type or ruvB::Tn26 as described above. Minocycline was administered intraperitoneally at doses of 0, 0.39, 0.78, 1.56, 3.12, 6.24, and 12.48 mg/kg 1 h post-infection, followed by a second dose at 24 h post-infection. Mice were euthanized at 30 h post-infection, and lungs, liver, spleen, and kidney were harvested for bacterial burden quantification(27). Homogenates were serially diluted and plated for bacterial enumeration, and bacterial burdens were normalized to tissue weight and expressed as CFU g⁻¹ tissue. For samples with no recoverable colonies, sample-specific limits of detection were calculated from the plated volume, homogenization volume and tissue weight. Data represent individual animals.

### Statistical Analysis

Statistical analyses were performed using GraphPad Prism v10. Biofilm biomass and confocal biofilm-thickness measurements were analyzed using ordinary one-way ANOVA followed by the indicated multiple-comparisons test. Growth curves were quantified by calculating the area under the curve (AUC) for each biological replicate and analyzed using ordinary one-way ANOVA with Šídák’s multiple-comparisons test. DNase I biofilm experiments were analyzed using two-way ANOVA with Tukey’s multiple-comparisons test. Pulmonary bacterial burden between two groups was compared using an unpaired two-tailed *t*-test with Welch’s correction. Survival distributions were compared using the log-rank (Mantel-Cox) test. Bacterial dissemination to the liver, spleen and kidney, as well as the minocycline dose-response experiments, were analyzed using two-way ANOVA with Šídák’s multiple-comparisons test. MIC assays were performed in three independent biological experiments; because identical MIC endpoints were obtained across replicates, no inferential statistical analysis was applied to MIC values. Unless otherwise stated, data are presented as mean ± s.d., and *P* < 0.05 was considered statistically significant. Exact *P* values and sample sizes are reported in the corresponding figure legends.

## Results

### AB5075 validated as representative model for tetracycline susceptibility studies

We first assessed whether AB5075, the genetic background of the Manoil transposon library, was representative of other A. baumannii isolates with respect to tetracycline-class susceptibility. MICs were measured for minocycline (MIN), doxycycline (DOX), tetracycline (TET), and tigecycline (TGC), across AB5075, ATCC 19606, and three additional clinical isolates (AB1605, AB1789, AB1710) (Fig. 1a). MIC values were reproducible and identical across three biological replicates.

**Fig 1.**
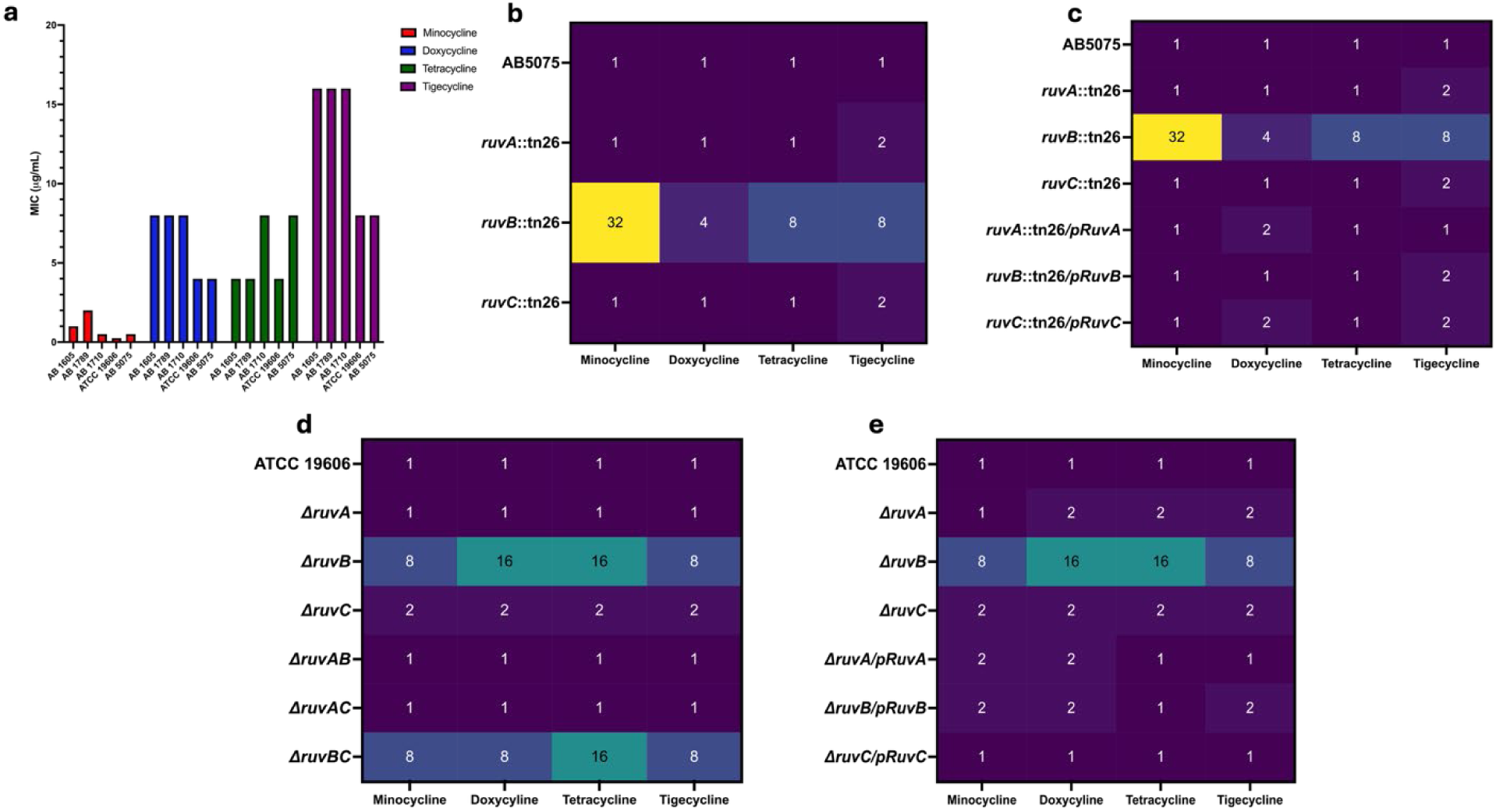
RuvB disruption increases tetracycline resistance in Acinetobacter baumannii. a, Minimum inhibitory concentrations (MICs) of minocycline (MIN), doxycycline (DOX), tetracycline (TET) and tigecycline (TGC) across five A. baumannii strains. b, Tetracycline-class MICs of AB5075 WT and ruvA::Tn26, ruvB::Tn26 and ruvC::Tn26 mutants. c, MICs of AB5075 Ruv mutants and their corresponding complemented strains. d, MICs of ATCC 19606 WT and single (ΔruvA, ΔruvB and ΔruvC) and double (ΔruvAB, ΔruvAC and ΔruvBC) Ruv deletion mutants. e, MICs of ATCC 19606 single Ruv deletion mutants and their corresponding complemented strains. MIC fold changes were calculated relative to the corresponding parental WT. Data are from three independent biological experiments (n = 3), with identical MIC endpoints obtained across replicates.

AB5075 displayed MICs of 0.5 µg/mL MIN, 8 µg/mL TET and TGC, and 4 µg/mL DOX, closely matching values observed for the comparator strains (AB1605: 1/8/4/16 µg/mL; AB1789: 2/8/4/16 µg/mL; AB1710: 0.5/8/8/16 µg/mL; ATCC 19606: 0.25/4/4/8 µg/mL). These results, together with prior reports establishing AB5075 as a widely used, genetically tractable MDR model strain and the availability of a curated transposon library, support using AB5075 and the Manoil library to dissect tetracycline responses and biofilm phenotypes(12, 13). We further used ATCC 19606 to see if the phenotype shown by AB5075 is strain dependent or carries through different strains of *A. baumannii*.

### RuvB disruption confers high-level resistance in AB5075 and ATCC 19606

#### AB5075 transposon mutants

In AB5075, disruption of ruvB::Tn26 caused a striking increase in minocycline resistance: the MIC rose from 0.5 µg/mL in WT to 16 µg/mL (32-fold) (Fig 1b.). This effect extended to other tetracyclines, with doxycycline increasing from 4 µg/mL (WT) to 16 µg/mL (4-fold)., tigecycline and tetracycline MICs each increasing from 8 µg/mL (WT) to 64 µg/mL (8-fold). By contrast, ruvA::Tn26 showed no effect on minocycline (0.5 µg/mL) and only a modest change in tigecycline (2-fold), while ruvC::Tn26 also produced limited shifts (TIG 2-fold). Complementation of each mutant restored MICs to near-WT values (Fig 1c.). MIC values were identical across three independent biological replicates.

#### ATCC 19606 markerless mutants

In ATCC 19606, deletion of ruvB raised the MIC for minocycline from 0.25 µg/mL (WT) to 2 µg/mL (8-fold), with parallel increases in doxycycline (4 → 64 µg/mL, 16-fold), tetracycline (4 → 64 µg/mL, 16-fold), and tigecycline (8 → 64 µg/mL, 8-fold) (Fig 1d.). ΔruvA had no effect (MIN 0.25 µg/mL), while ΔruvC produced only modest two fold increases on all tetracyclines (0.25 → 0.5 µg/mL, 2-fold). Double mutants separated these contributions: ΔruvAB resembled WT across all antibiotics, consistent with the hypothesis that free RuvA liberated in the absence of RuvB is responsible for the phenotype; ΔruvBC mirrored ΔruvB; ΔruvAC was unchanged. Complementation of single ruv deletion mutants restored MICs to WT values (Fig 1e.). These genetic dependencies align with the canonical RuvAB-RuvC mechanism for Holliday junction (HJ) processing, in which RuvA binds the junction, RuvB motors drive branch migration, and RuvC resolves the structure(28–30).

### Loss of ruvB drives robust biofilm formation in both AB5075 and ATCC 19606

Biofilm biomass was quantified by the crystal violet microtiter assay. In AB5075, ruvB::Tn26 mutants produced a ∼550% increase in biomass relative to WT (mean ± s.d., n = 6 biological replicates; ordinary one-way ANOVA with Tukey’s multiple comparisons test; P < 0.0001), whereas ruvA::Tn26 and ruvC::Tn26 showed no significant difference (P = 0.9930 and P = 0.9972, respectively) (Fig 2.). Complementation of ruvB reduced biomass back to WT levels (Fig 2a).

**Fig 2.**
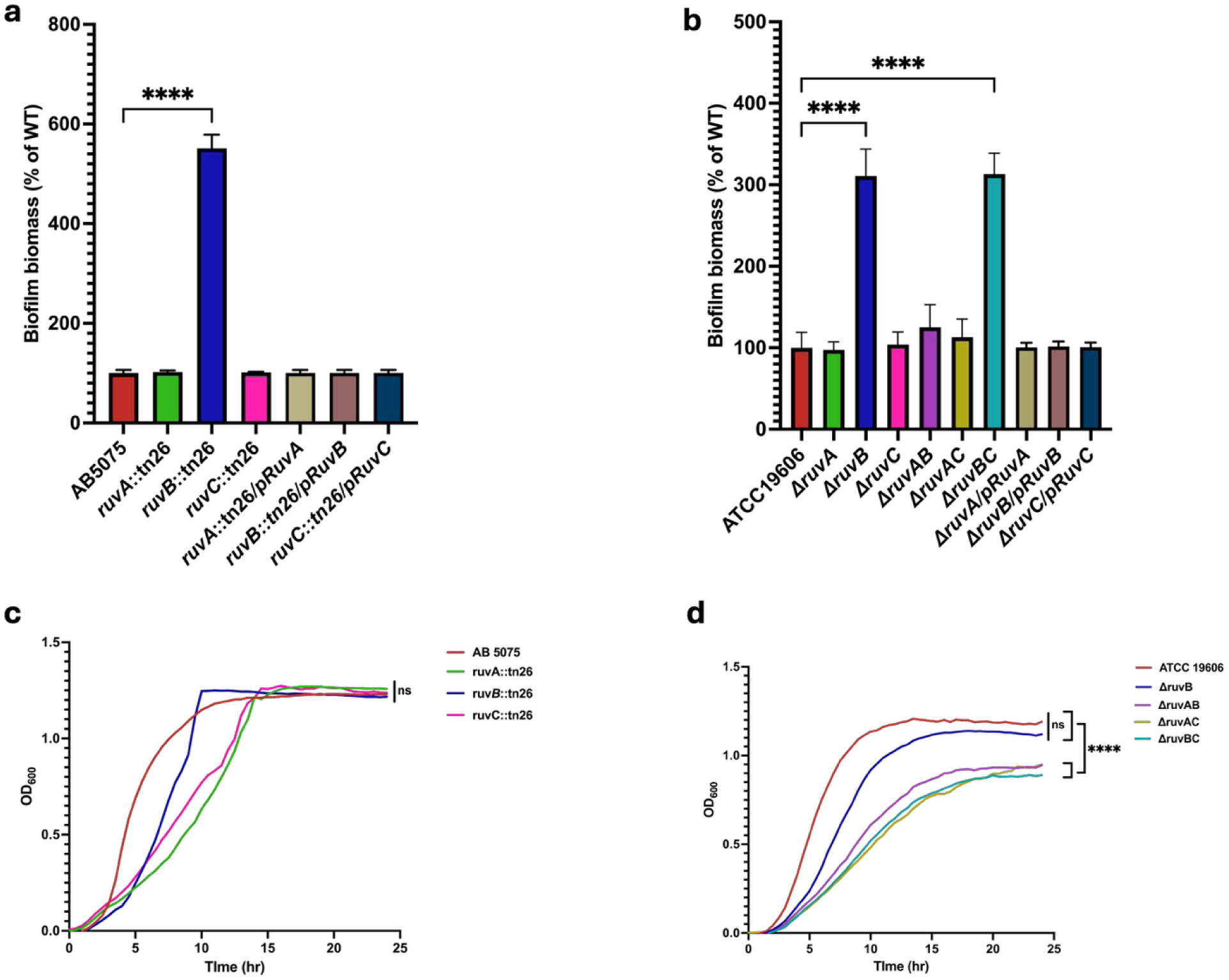
Loss of RuvB promotes biofilm formation independently of planktonic growth. a, Crystal violet biofilm biomass of AB5075 WT, Ruv transposon mutants and complemented strains, expressed relative to WT. Data are mean ± s.d. from six biological replicates. Statistical significance was assessed by ordinary one-way ANOVA with Tukey’s multiple-comparisons test; ruvB::Tn26 versus WT, P < 0.0001; ruvA::Tn26 versus WT, P = 0.9930; ruvC::Tn26 versus WT, P = 0.9972. b, Crystal violet biofilm biomass of ATCC 19606 WT, single and double Ruv deletion mutants and complemented strains, expressed relative to WT. Data are mean ± s.d. from eight biological replicates. Statistical significance was assessed by ordinary one-way ANOVA with Tukey’s multiple-comparisons test; ΔruvB versus WT, P < 0.0001; ΔruvA versus WT, P > 0.9999; ΔruvC versus WT, P = 0.9998; ΔruvAB versus WT, P = 0.3143; ΔruvBC versus WT, P < 0.0001. c, Growth kinetics of AB5075 WT and ruv::Tn26 in CA-MHB without antibiotics. Growth was quantified by area under the curve (AUC) from three biological replicates and compared using ordinary one-way ANOVA with Šídák’s multiple-comparisons test; WT versus ruvA::Tn26, P = 0.1786; WT versus ruvB::Tn26, P = 0.8463; WT versus ruvC::Tn26, P = 0.3862. d, Growth kinetics of ATCC 19606 WT, ΔruvB, ΔruvAB, ΔruvAC and ΔruvBC in CA-MHB without antibiotics. AUC values were analyzed from three biological replicates using ordinary one-way ANOVA with Šídák’s multiple-comparisons test; WT versus ΔruvB, P = 0.3345; WT versus ΔruvAB, P = 0.0005; WT versus ΔruvAC and ΔruvBC, P < 0.0001.

In ATCC 19606, ΔruvB mutants showed a ∼300% biomass increase relative to wild type (mean ± s.d., n = 8 biological replicates; ordinary one-way ANOVA with Tukey’s multiple comparisons test; P < 0.0001) (Fig 2b.). Deletion of ruvA or ruvC did not significantly alter biomass (P > 0.9999 and P = 0.9998 respectively). Double-mutant analysis revealed that ΔruvAB resembled WT (P=0.3143), whereas ΔruvBC phenocopied ΔruvB (P < 0.0001), reinforcing the role of free RuvA. Complementation normalized biomass in both cases (Fig 2b.). Assay execution followed community standards(21).

### Growth kinetics demonstrate comparable planktonic growth across WT and ruv mutants

To verify that increased biofilm biomass in ruv mutants was not an artifact of enhanced planktonic growth, we measured growth kinetics in cation-adjusted Mueller-Hinton broth (CA-MHB) without antibiotics (n = 3 biological replicates). Growth curves were quantified by area under the curve (AUC) and analyzed using ordinary one-way ANOVA with Šídák’s multiple comparisons test (Fig 2.).

In AB5075, ruvB::Tn26 displayed growth curves indistinguishable from WT, with overlapping lag, exponential, and stationary phases (P = 0.8463). Similarly, in ATCC 19606, ΔruvB exhibited growth kinetics comparable to WT based on AUC analysis (P = 0.3345). In contrast, ATCC 19606 ΔruvAB, ΔruvAC and ΔruvBC strains exhibited reduced growth relative to wild type (P = 0.0005, P < 0.0001, and P < 0.0001, respectively). These findings confirm that the exaggerated biofilm phenotypes of ruvB-deficient strains are not attributable to increased cell proliferation but instead reflect true increases in extracellular matrix content, particularly eDNA.

### Confocal microscopy reveals thicker eDNA-rich biofilms in ruvB mutants

Confocal Z-stacks revealed marked structural changes in ruvB biofilms. Biofilms were stained using a four-channel fluorescent panel to simultaneously visualize cells, proteins, lipids, DNA, and extracellular DNA, including SYTOX Red as a high-specificity marker for extracellular DNA within the biofilm matrix. In AB5075, biomass increased ∼260% (P < 0.001), and mean thickness rose from 5.56 µm (WT) to 46.00 µm in ruvB::Tn26 (Fig. 3 a-b). ruvA::tn26 mutants formed thinner biofilms (7.84 µm), while ruvC::tn26 had 11.55 µm. Complementation of ruvB restored biofilm thickness to WT levels. In ATCC 19606, ΔruvB mutants formed thicker biofilms (49.1 µm vs 5.56 µm WT, P < 0.0001), ΔruvBC mirrored ΔruvB (45.2 µm), and ΔruvAB had a thickness of 11.5 µm.

**Fig 3.**
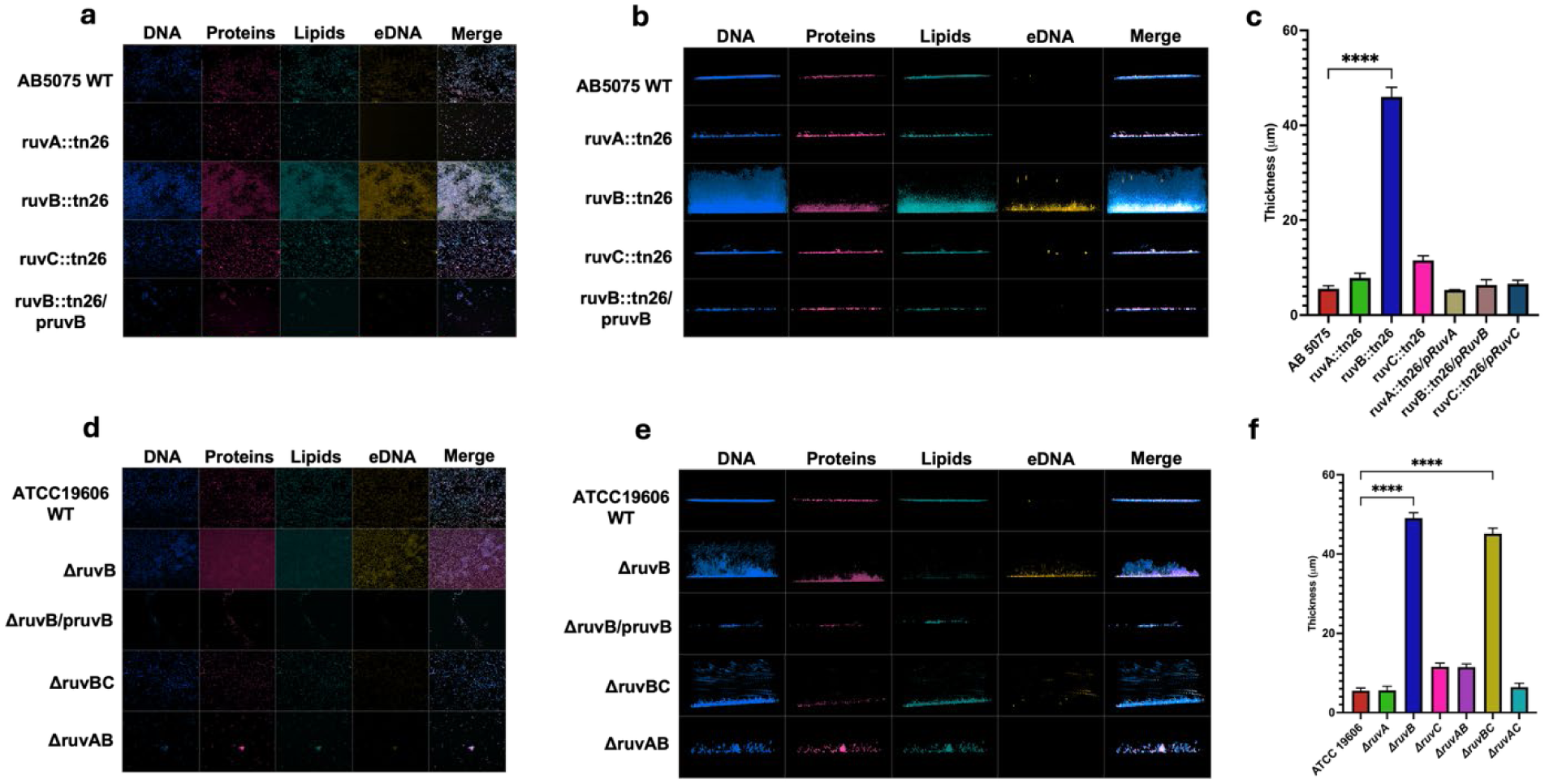
RuvB-deficient biofilms are thicker and enriched in extracellular DNA. a, Representative confocal images of 72-h AB5075 WT, Ruv transposon mutant and complemented biofilms stained with Hoechst 33258 for DNA, SYTOX Red for extracellular DNA, BODIPY FL NHS ester for lipids and Texas Red hydrazide for proteins. b, Orthogonal views of the corresponding confocal Z-stacks showing biofilm vertical architecture. c, Quantification of AB5075 biofilm thickness. Data are mean ± s.d. from three biological replicates (n = 3). Statistical significance was assessed by ordinary one-way ANOVA with Šídák’s multiple-comparisons test; ruvB::Tn26 versus WT, P < 0.0001. d, Representative confocal images of 72-h ATCC 19606 WT and Ruv deletion mutant biofilms stained as in a. e, Orthogonal views of the corresponding ATCC 19606 confocal Z-stacks. f, Quantification of ATCC 19606 biofilm thickness. Data are mean ± s.d. from (n = 3). Statistical significance was assessed by ordinary one-way ANOVA with Šídák’s multiple-comparisons test; ΔruvB versus WT, P < 0.0001. Three-dimensional reconstructions of representative biofilms are provided in Supplementary Videos 1 and 2.

### DNase I treatment revealss an eDNA-dependent mechanism in ruvB mutants

To assess the contribution of extracellular DNA to biofilm integrity, established biofilms were treated with DNase I (5 µg/mL) for 24 hours (n = 6 biological replicates). Biofilm biomass was normalized to the untreated WT condition, which was defined as 100%. Statistical analysis was performed using two-way ANOVA with Tukey’s multiple comparisons test. (Fig 4)

**Fig 4.**
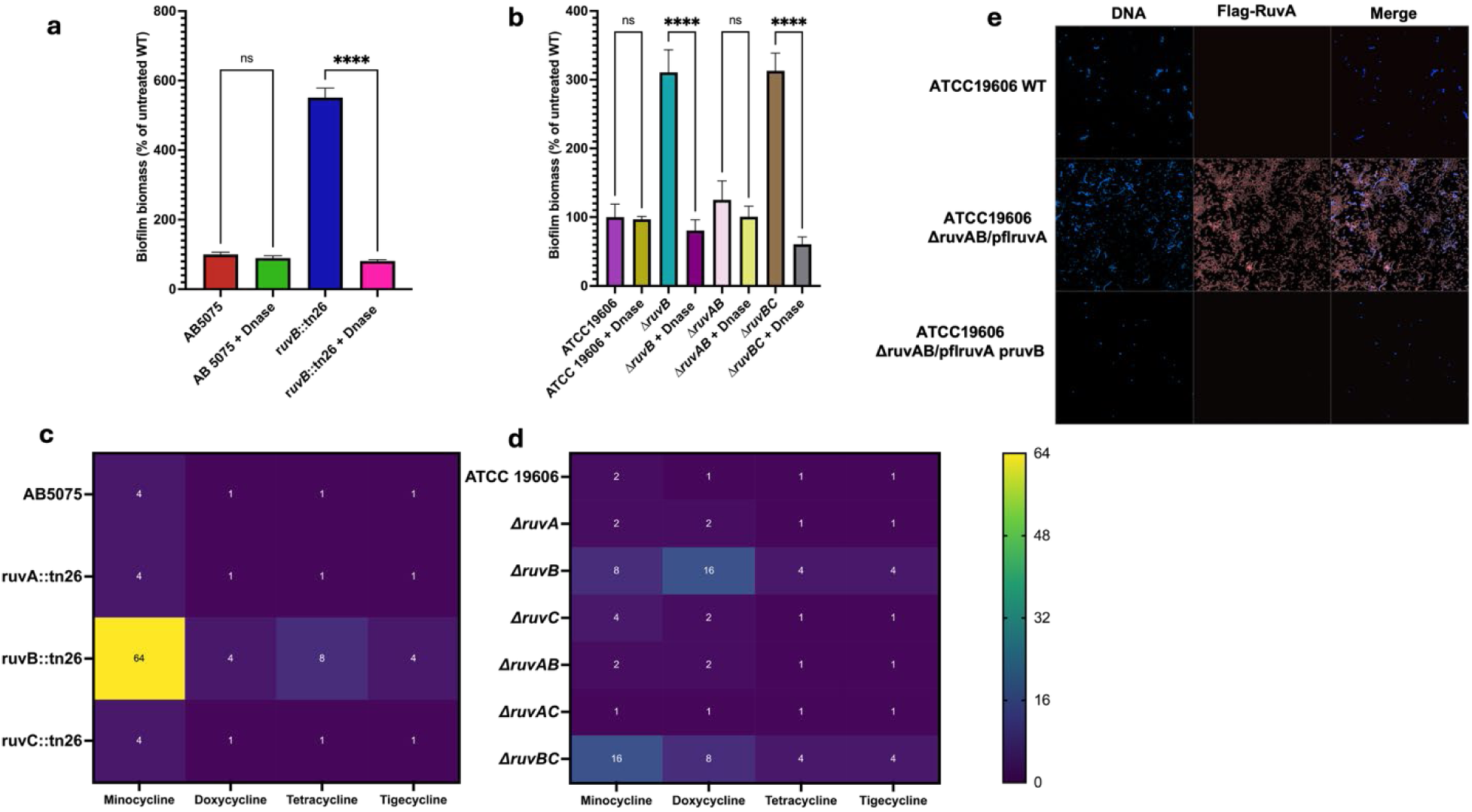
Extracellular DNA contributes to RuvB-dependent biofilm formation and tetracycline resistance. **a,** Biofilm biomass of AB5075 WT and *ruvB*::Tn26 following DNase I treatment, normalized to untreated WT, which was defined as 100%. WT untreated versus DNase I-treated, *P* = 0.9512; *ruvB*::Tn26 untreated versus DNase I-treated, *P* < 0.0001. **b,** Biofilm biomass of ATCC 19606 WT and indicated *ruv* deletion mutants following DNase I treatment, normalized to untreated WT, which was defined as 100%. WT untreated versus DNase I-treated, *P* > 0.9999; Δ*ruvB*, *P* < 0.0001; Δ*ruvAB*, *P* = 0.3030; Δ*ruvBC*, *P* < 0.0001. Data in **a,b** are mean ± s.d. from six biological replicates (n = 6). Statistical significance was assessed by two-way ANOVA with Tukey’s multiple-comparisons test. **c,d,** DNase I-mediated fold reduction in the MICs of minocycline, doxycycline, tetracycline and tigecycline for AB5075 WT and Ruv transposon mutants (**c**) and ATCC 19606 WT and indicated *ruv* deletion mutants (**d**). Fold reduction was calculated as the MIC in the absence of DNase I divided by the MIC in the presence of DNase I; values of 1 indicate no change in MIC. MIC assays were performed in three independent biological experiments (n = 3), with identical MIC endpoints across replicates. **e,** Representative confocal images of ATCC 19606 WT, Δ*ruvAB* expressing FLAG–RuvA, and Δ*ruvAB* expressing FLAG-RuvA together with complemented *ruvB*. DNA was visualized with Hoechst 33258 and FLAG–RuvA was detected by anti-FLAG immunofluorescence.

In AB5075, DNase I treatment had no significant effect on wild-type biofilms (P = 0.9512), whereas ruvB::Tn26 biofilms exhibited ∼85-90% reduction in biomass relative to untreated controls (P < 0.0001). Similarly, in ATCC 19606, DNase treatment did not alter wild-type biomass (P > 0.9999), while ΔruvB and ΔruvBC biofilms were ∼85-90% reduced following DNase exposure (P < 0.0001 for both comparisons). In contrast, ΔruvAB biofilms were not significantly affected by DNase treatment (P = 0.3030).

These data, together with the classic and contemporary literature on eDNA-dependent biofilm integrity, support an eDNA stabilized matrix in ruvB deficient settings(31–34). The specific dose of DNase I we used is consistent with published efficacious ranges (35, 36).

### DNase I addition reverts tetracycline resistance in ruvB mutants and reduces MICs in wild-type strains

To directly test whether extracellular DNA stabilization underpins the elevated MICs of ruvB mutants, we repeated tetracycline-class MIC assays in the presence of 5 µg/mL DNase I throughout the incubation period (n = 3 biological replicates). MIC values were identical across independent experiments. In AB5075, DNase I markedly reduced the elevated tetracycline-class MICs of ruvB::Tn26, including a 64-fold reduction in minocycline MIC from 16 to 0.25 µg/mL. In ATCC 19606, DNase I also markedly reduced tetracycline-class MICs in ΔruvB and ΔruvBC mutants (e.g., MIN 2 → 0.25 µg/mL, DOX 64 → 4 µg/mL, TET 64 → 16 µg/mL, TGC 64 → 16 µg/mL) (Fig 4.). Notably, DNase I also reduced selected MICs in the parental strains, most notably a fourfold reduction in the AB5075 minocycline MIC. These results demonstrate that eDNA makes a major contribution to the elevated tetracycline resistance associated with ruvB loss.

### FLAG-tagged RuvA accumulates within the biofilm matrix in the absence of RuvB

To determine whether RuvA accumulates within the biofilm matrix in the absence of RuvB, we visualized FLAG-tagged RuvA using confocal microscopy. A ruvAB double mutant was complemented with a FLAG-tagged ruvA allele (kanamycin-resistant), allowing specific detection of RuvA independently of native RuvB activity. Biofilms were stained with Hoechst to visualize DNA and probed with an anti-FLAG (DYKDDDDK) monoclonal antibody conjugated to Alexa Fluor™ Plus 594 to detect FLAG-tagged RuvA. Robust FLAG signal was observed exclusively in the ΔruvAB strain complemented with FLAG-tagged ruvA, where FLAG-RuvA-associated fluorescence accumulated prominently within the biofilm matrix. In contrast, no FLAG signal was detected in wild-type strains or in ruvAB mutants fully complemented with both FLAG-tagged ruvA and wild-type ruvB, despite identical staining and imaging conditions (Fig 4e.). Restoration of ruvB abolished extracellular localization of FLAG-tagged RuvA, indicating that RuvB suppresses RuvA accumulation in the biofilm matrix. Together with the eDNA enrichment and DNase-sensitive phenotype of RuvB-deficient biofilms, these findings support an association between RuvA accumulation and the eDNA-rich biofilm matrix.

### RuvB disruption results in comparable virulence and pulmonary bacterial burden in a murine pneumonia model

To determine whether ruvB-dependent biofilm and resistance phenotypes translated to altered in vivo infection outcomes, we evaluated virulence of AB5075 wild-type and ruvB mutant strains using a murine pneumonia model (n = 5 mice per group). Survival was monitored over six days and analyzed using the log-rank (Mantel-Cox) test. All mice in both groups succumbed during the observation period, with no significant difference in survival between groups (log-rank Mantel– Cox test, P = 0.4945). Pulmonary bacterial burden was quantified at the time of euthanasia by enumerating bacterial burden as CFU g⁻¹ lung tissue. Consistent with the survival data, lung bacterial burden analysis demonstrated comparable pulmonary CFU between wild-type and ruvB mutant infected mice at the time of euthanasia (unpaired t-test with Welch’s correction; P = 0.08). These findings indicate that ruvB disruption results in comparable virulence or bacterial replication in the lung during pneumonia.

To assess bacterial dissemination beyond the lungs, bacterial burdens were quantified in the liver, spleen, and kidneys at the time of euthanasia. Wild-type infections exhibited higher bacterial loads across all secondary organs compared to ruvB::Tn26-infected mice, whereas ruvB::Tn26 showed reduced CFU, indicating diminished dissemination from the primary site of infection. Bacterial burdens were significantly lower in ruvB::Tn26-infected mice in the liver, spleen and kidney (two-way ANOVA with Šídák’s multiple-comparisons test; P < 0.0001 for each organ; Fig. 5c)

**Fig 5.**
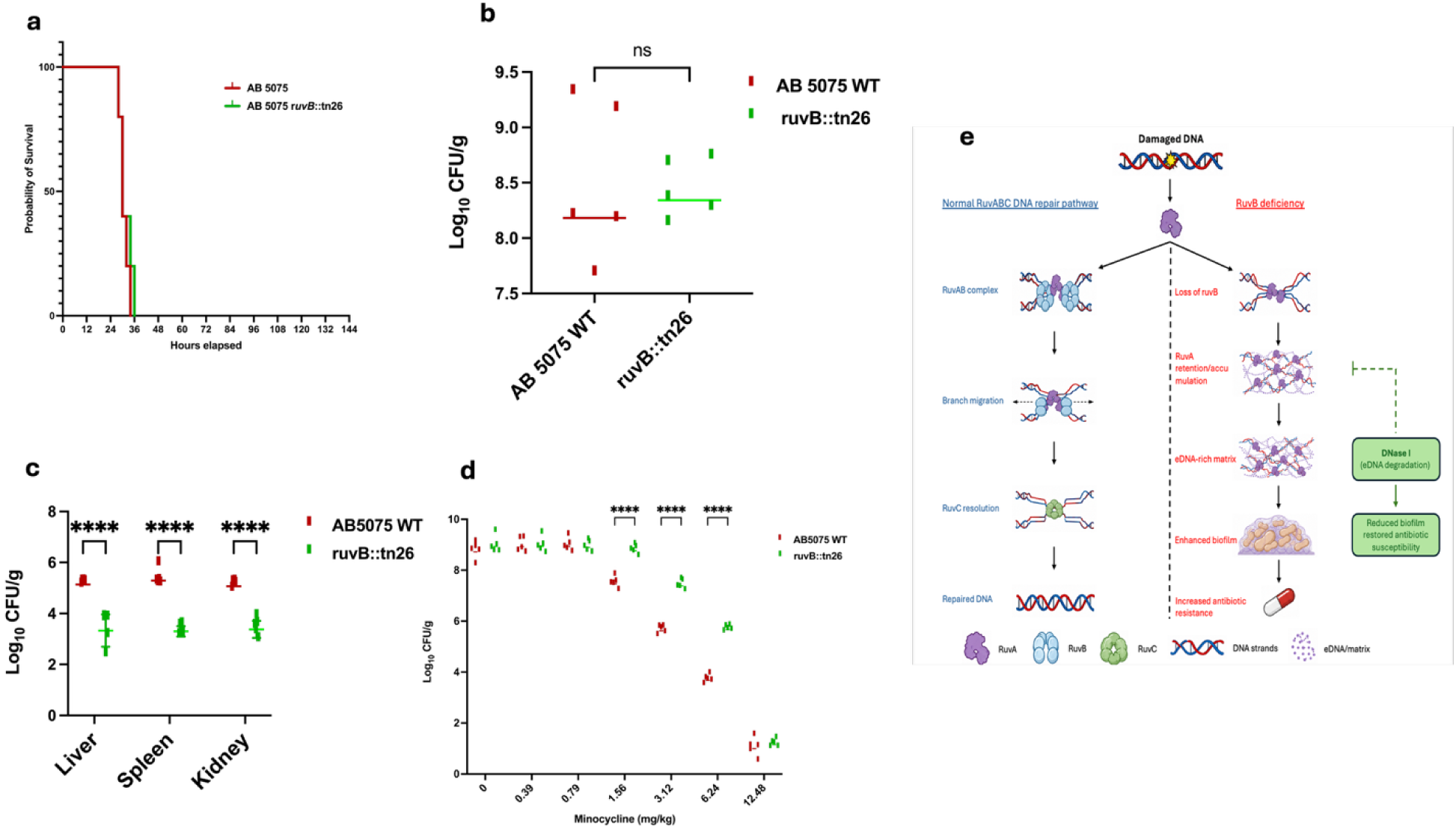
RuvB disruption alters bacterial dissemination and minocycline responsiveness during acute pneumonia. a, Kaplan-Meier survival of C57BL/6 mice infected with AB5075 WT or ruvB::Tn26 (n = 5 mice per group). Survival was compared using a log-rank (Mantel–Cox) test; P = 0.4945. b, Pulmonary bacterial burden of AB5075 WT and ruvB::Tn26 at 30 h after infection. Each point represents one mouse, and horizontal lines indicate the mean (n = 5 mice per group). Data are expressed as log₁₀(CFU g⁻¹ lung tissue). Groups were compared using an unpaired two-tailed t-test with Welch’s correction; P = 0.08. c, Bacterial burden in the liver, spleen and kidney following infection with AB5075 WT or ruvB::Tn26. Data are expressed as log₁₀(CFU g⁻¹ tissue); each point represents one mouse, and horizontal lines indicate the mean (n = 5 mice per group). Statistical significance was assessed by two-way ANOVA with Šídák’s multiple-comparisons test; liver, spleen and kidney, P < 0.0001. Samples with no recoverable CFU were plotted at the sample-specific limit of detection. d, Pulmonary bacterial burden following minocycline treatment at 0, 0.39, 0.78, 1.56, 3.12, 6.24 and 12.48 mg kg⁻¹ in mice infected with AB5075 WT or ruvB::Tn26. Each point represents one mouse, and horizontal lines indicate the mean (n = 5 mice per group). Data are expressed as log₁₀(CFU g⁻¹ lung tissue). Statistical significance was assessed by two-way ANOVA with Šídák’s multiple-comparisons test; WT versus ruvB::Tn26, P < 0.0001 at doses 1.56, 3.12, and 6.24 mg kg⁻¹; other comparisons were not significant. e, working model of RuvB-dependent regulation of RuvA-associated extracellular DNA, biofilm formation and antibiotic resistance.

### Minocycline treatment reveals reduced antibiotic responsiveness in ruvB-deficient infections

To determine whether ruvB-dependent biofilm phenotypes influence antibiotic treatment outcomes in vivo, we performed a minocycline dose response study in a murine pneumonia model. C57BL/6 mice were intratracheally infected with AB5075 wild type or ruvB::Tn26 and treated with increasing doses of minocycline (0, 0.39, 0.78, 1.56, 3.12, 6.24, 12.48 mg/kg) administered 1 h post-infection, followed by a second dose at 24 h. Lungs were harvested at 30 h post-infection for bacterial burden quantification (n = 5 mice per group).In wild-type infections, minocycline treatment resulted in a dose-dependent reduction in lung bacterial burden, with progressively lower CFU observed at increasing doses (Fig 5d.). In contrast, ruvB::Tn26-infected mice exhibited attenuated reduction in bacterial burden across the same dosing range, with consistently higher residual CFU compared to wild type, particularly at intermediate treatment doses. Statistical analysis using two-way ANOVA with Šídák’s multiple-comparisons test identified significantly higher pulmonary bacterial burdens in ruvB::Tn26-infected mice compared with WT-infected mice at 1.56, 3.12 and 6.24 mg kg⁻¹ minocycline (P < 0.0001 for each comparison), whereas the remaining dose comparisons were not significant (Fig. 5d). These data indicate that ruvB disruption is associated with reduced responsiveness to minocycline treatment in vivo, particularly at higher antibiotic exposure, consistent with increased biofilm-associated resistance observed in vitro.

### Free RuvA stabilizes eDNA lattices, providing a unifying mechanism for biofilm-associated resistance

Integrating genetic and imaging data, we propose that loss of RuvB liberates RuvA from the branch-migration motor, enabling free RuvA to bind HJ-like junctions within the eDNA lattice of biofilms. This stabilizes the matrix, increases biofilm thickness, and elevates antibiotic resistance (Fig. 5e). The model is consistent with structural and mechanistic work on RuvAB-HJ processing and the observation that biofilm eDNA forms HJ-related lattices (11, 28–30). To our knowledge, this is the first demonstration that a DNA repair protein complex can directly stabilize extracellular DNA to drive biofilm-associated resistance.

## Discussion

Biofilm-associated antibiotic resistance is a defining feature of Acinetobacter baumannii infections, yet the molecular mechanisms that couple intracellular stress responses to extracellular matrix organization remain incompletely understood(37–39). Here, we identify the Holliday junction branch migration motor RuvB as a key regulator of extracellular DNA (eDNA) dependent biofilm architecture and tetracycline resistance. By combining genetic dissection, quantitative imaging, enzymatic perturbation, and in vivo infection models, we show that loss of RuvB uncouples canonical DNA repair from matrix control, allowing RuvA to stabilize eDNA lattices and promote antibiotic tolerance.

Disruption of ruvB conferred marked resistance to tetracycline-class antibiotics in two genetically distinct *A. baumannii* backgrounds, whereas deletion of ruvA abolished this phenotype and deletion of ruvC had only modest effects. Double-mutant analysis further demonstrated that resistance depends on the presence of RuvA in the absence of RuvB, consistent with the established roles of RuvA as a Holliday junction-binding protein and RuvB as its cognate branch-migration motor (40–42). These genetic dependencies support a model in which the resistance phenotype is driven, at least in part, by RuvA-dependent matrix effects arising in the absence of RuvB, rather than by loss of DNA repair alone.

Multiple assays demonstrated that ruvB-dependent resistance is associated with altered biofilm matrix structure rather than increased bacterial growth. ruvB mutants formed substantially thicker biofilms with increased biomass despite indistinguishable planktonic growth kinetics. High-resolution confocal microscopy using complementary DNA stains, including SYTOX Red to selectively label extracellular DNA, revealed dense, lattice-like eDNA networks within ruvB-deficient biofilms. These data establish that the observed phenotypes reflect restructuring of the extracellular matrix rather than increased cell number or intracellular DNA release.

Direct visualization of FLAG-tagged RuvA provides mechanistic insight into how these eDNA structures are stabilized. In ruvAB mutants complemented with FLAG-tagged ruvA, FLAG-RuvA accumulated prominently within the biofilm matrix in the absence of RuvB, whereas restoration of RuvB markedly reduced this signal. This localization was absent in wild-type strains and in ruvAB mutants complemented with both ruvA and ruvB. The marked reduction in FLAG-RuvA-associated signal following restoration of RuvB is consistent with RuvB limiting matrix-associated accumulation of RuvA.

Functional perturbation experiments further support a central role for eDNA stabilization in ruvB-dependent antibiotic resistance. DNase I treatment collapsed ruvB mutant biofilms and markedly reduced elevated tetracycline-class MICs, supporting a major contribution of eDNA to the resistance phenotype. The modest reduction in MICs observed in wild-type strains upon DNase treatment suggests that basal eDNA contributes to intrinsic antibiotic resistance even in the absence of genetic perturbation, consistent with previous work implicating eDNA as a structural and protective component of bacterial biofilms(31, 36, 43). Together with SYTOX Red enrichment and the DNase-sensitive phenotype, these observations support an association between RuvA accumulation and the eDNA-rich matrix.

Despite these pronounced in vitro phenotypes, ruvB disruption resulted in comparable mortality and pulmonary bacterial burden to wild-type AB5075 in a murine pneumonia model. To further assess the relevance of this phenotype under therapeutic conditions, we performed a minocycline dose response study in vivo. While wild-type infections exhibited a reduction in bacterial burden with increasing antibiotic dose, ruvB-deficient infections displayed diminished responsiveness to treatment, with significantly higher lung burdens in ruvB::Tn26-infected mice at 1.56, 3.12 and 6.24 mg kg⁻¹, whereas the groups did not differ significantly at 12.48 mg kg⁻¹. Notably, ruvB-deficient infections exhibited reduced dissemination to secondary organs, suggesting that while acute pulmonary infection remains unchanged, alterations in extracellular DNA dynamics may influence bacterial spread within the host. These findings also indicate that ruvB-dependent extracellular DNA stabilization limits antibiotic efficacy under treatment conditions. This dissociation between biofilm-associated antibiotic resistance and virulence highlights the context-specific nature of eDNA-mediated phenotypes. Acute pneumonia models predominantly capture planktonic growth and host damage, whereas biofilm-mediated resistance is likely to play a greater role during chronic infection or antibiotic exposure(44–46).

Together, these findings support a model in which RuvB functions as a gatekeeper that limits RuvA activity to intracellular Holliday junction processing. Loss of RuvB liberates RuvA, promoting RuvA accumulation within the biofilm matrix, consistent with a model in which RuvA contributes to stabilization of eDNA-rich structures. This work reveals an unexpected connection between DNA repair machinery and extracellular matrix organization and identifies eDNA-protein interactions as a potential target for disrupting biofilm-associated antibiotic tolerance without directly impairing bacterial viability(11, 47, 48).

### Conclusion

In summary, this study identifies the DNA repair motor RuvB as a key regulator of extracellular DNA dependent biofilm architecture and antibiotic resistance in *Acinetobacter baumannii*. We show that loss of RuvB liberates RuvA from its canonical intracellular role, enabling direct stabilization of extracellular DNA lattices within the biofilm matrix. This eDNA-RuvA interaction promotes biofilm thickening and tetracycline resistance without measurably affecting outcomes in an acute pneumonia model, highlighting a functional distinction between antibiotic tolerance and host lethality. By linking Holliday junction processing machinery to extracellular matrix organization, our findings reveal an unexpected mechanism by which genome maintenance proteins can shape multicellular bacterial behaviors. More broadly, this work highlights eDNA-protein interactions as tractable targets for disrupting biofilm associated antibiotic resistance without directly impairing bacterial growth.

---

**Table S1:**
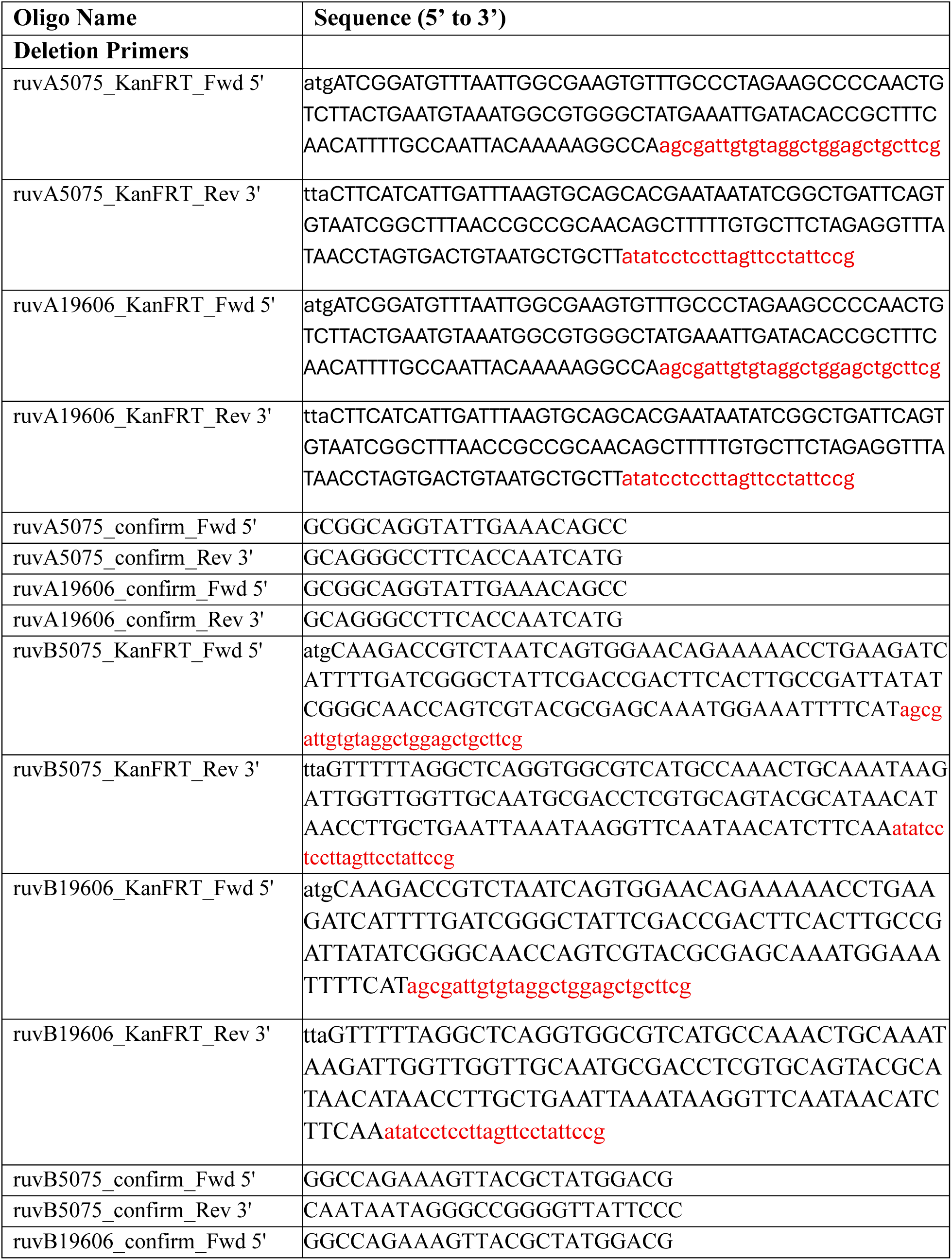
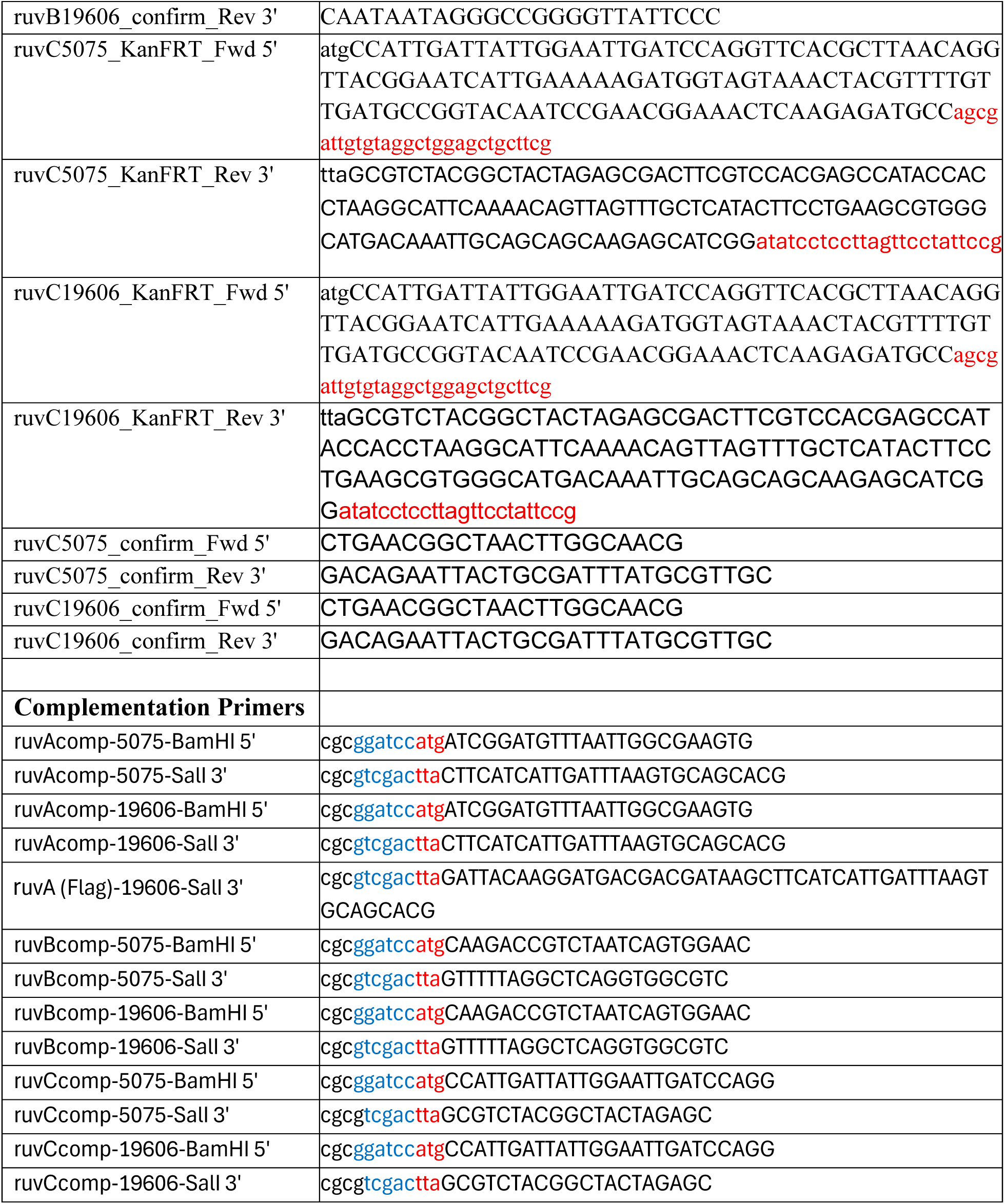
Primers used in this study.

## Reference

1. Murray CJL, Ikuta KS, Sharara F, Swetschinski L, Robles Aguilar G, Gray A, et al. Global burden of bacterial antimicrobial resistance in 2019: a systematic analysis. The Lancet. 2022;399(10325):629–55.

2. Sharma R, Lakhanpal D. Acinetobacter baumannii: A comprehensive review of global epidemiology, clinical implications, host interactions, mechanisms of antimicrobial resistance and mitigation strategies. Microb Pathog. 2025;204:107605.

3. Goncalves EK, Penwell WF, Fiester SE. Combating the Growing Threat of Acinetobacter baumannii Resistance. Antibiotics. 2025;14(7):694.

4. Tsakris A, Koumaki V, Dokoumetzidis A. Minocycline susceptibility breakpoints for Acinetobacter baumannii: do we need to re-evaluate them? Journal of Antimicrobial Chemotherapy. 2018;74(2):295–7.

5. Velmurugan P, Ramalingam AJ, Saikumar C. An Ancient Drug for a Modern Era: Minocycline for the Treatment of Multi-Drug-Resistant Acinetobacter baumannii. Cureus. 2024;16(6):e61785.

6. Roca Subirà I, Espinal P, Vila-Farrés X, Vila Estapé J. The Acinetobacter baumannii Oxymoron: Commensal Hospital Dweller Turned Pan-Drug-Resistant Menace. Frontiers in Microbiology. 2012;Volume 3 - 2012.

7. Harding CM, Tracy EN, Carruthers MD, Rather PN, Actis LA, Munson RS. Acinetobacter baumannii Strain M2 Produces Type IV Pili Which Play a Role in Natural Transformation and Twitching Motility but Not Surface-Associated Motility. mBio. 2013;4(4):10.1128/mbio.00360-13.

8. Foster PL. Stress-Induced Mutagenesis in Bacteria. Critical Reviews in Biochemistry and Molecular Biology. 2007;42(5):373–97.

9. Cirz RT, Chin JK, Andes DR, de Crécy-Lagard V, Craig WA, Romesberg FE. Inhibition of Mutation and Combating the Evolution of Antibiotic Resistance. PLOS Biology. 2005;3(6):e176.

10. Kohanski MA, Dwyer DJ, Collins JJ. How antibiotics kill bacteria: from targets to networks. Nature Reviews Microbiology. 2010;8(6):423–35.

11. Devaraj A, Buzzo JR, Mashburn-Warren L, Gloag ES, Novotny LA, Stoodley P, et al. The extracellular DNA lattice of bacterial biofilms is structurally related to Holliday junction recombination intermediates. Proceedings of the National Academy of Sciences. 2019;116(50):25068–77.

12. Jacobs AC, Thompson MG, Black CC, Kessler JL, Clark LP, McQueary CN, et al. AB5075, a Highly Virulent Isolate of Acinetobacter baumannii, as a Model Strain for the Evaluation of Pathogenesis and Antimicrobial Treatments. mBio. 2014;5(3):e01076-14.

13. Gallagher LA, Ramage E, Weiss EJ, Radey M, Hayden HS, Held KG, et al. Resources for Genetic and Genomic Analysis of Emerging Pathogen Acinetobacter baumannii. J Bacteriol. 2015;197(12):2027–35.

14. Tucker AT, Nowicki EM, Boll JM, Knauf GA, Burdis NC, Trent MS, et al. Defining gene-phenotype relationships in Acinetobacter baumannii through one-step chromosomal gene inactivation. mBio. 2014;5(4):e01313–14.

15. Oh MH, Lee JC, Kim J, Choi CH, Han K. Simple Method for Markerless Gene Deletion in Multidrug-Resistant Acinetobacter baumannii. Appl Environ Microbiol. 2015;81(10):3357–68.

16. Institute CaLS. CLSI vs FDA Breakpoints, 34th ed. CLSI Guideline M100. Clinical and Laboratory Standards Institute; 2024.

17. CLSI. Methods for Dilution Antimicrobial Susceptibility Tests for Bacteria That Grow Aerobically. 11th ed. CLSI standard M07. Wayne, PA: Clinical and Laboratory Standards Institute. 2018.

18. Olea-Ozuna RJ, Campbell MJ, Quintanilla SY, Nandy S, Brodbelt JS, Boll JM. Alternative lipid synthesis in response to phosphate limitation promotes antibiotic tolerance in Gram-negative ESKAPE pathogens. PLoS Pathog. 2025;21(2):e1012933.

19. Podolsky T, Fong ST, Lee BT. Direct selection of tetracycline-sensitive Escherichia coli cells using nickel salts. Plasmid. 1996;36(2):112–5.

20. Tiwari S, Nizet O, Dillon N. Development of a high-throughput minimum inhibitory concentration (HT-MIC) testing workflow. Frontiers in Microbiology. 2023;14.

21. O’Toole GA. Microtiter dish biofilm formation assay. J Vis Exp. 2011(47).

22. Hofer G, Steyrer E, Kostner GM, Hermetter A. LDL-mediated interaction of Lp[a] with HepG2 cells: a novel fluorescence microscopy approach. J Lipid Res. 1997;38(12):2411–21.

23. Hirose M, Tohda H, Giga-Hama Y, Tsushima R, Zako T, Iizuka R, et al. Interaction of a small heat shock protein of the fission yeast, Schizosaccharomyces pombe, with a denatured protein at elevated temperature. J Biol Chem. 2005;280(38):32586–93.

24. Titus JA, Haugland R, Sharrow SO, Segal DM. Texas Red, a hydrophilic, red-emitting fluorophore for use with fluorescein in dual parameter flow microfluorometric and fluorescence microscopic studies. J Immunol Methods. 1982;50(2):193–204.

25. Heller I, Sitters G, Broekmans OD, Farge G, Menges C, Wende W, et al. STED nanoscopy combined with optical tweezers reveals protein dynamics on densely covered DNA. Nature Methods. 2013;10(9):910–6.

26. Kamwouo T, Bouttier S, Domenichini S, Saunier J, Coullon H, Simons A, et al. Extracellular DNA filaments associated with surface polysaccharide II give Clostridioides difficile biofilm matrix a network-like structure. npj Biofilms and Microbiomes. 2025;11(1):108.

27. Dillon N, Holland M, Tsunemoto H, Hancock B, Cornax I, Pogliano J, et al. Surprising synergy of dual translation inhibition vs. Acinetobacter baumannii and other multidrug-resistant bacterial pathogens. EBioMedicine. 2019;46:193–201.

28. West SC. Processing of recombination intermediates by the RuvABC proteins. Annu Rev Genet. 1997;31:213–44.

29. Wald J, Fahrenkamp D, Goessweiner-Mohr N, Lugmayr W, Ciccarelli L, Vesper O, et al. Mechanism of AAA+ ATPase-mediated RuvAB-Holliday junction branch migration. Nature. 2022;609(7927):630–9.

30. Rish AD, Shen Z, Chen Z, Zhang N, Zheng Q, Fu T-M. Molecular mechanisms of Holliday junction branch migration catalyzed by an asymmetric RuvB hexamer. Nature Communications. 2023;14(1):3549.

31. Whitchurch CB, Tolker-Nielsen T, Ragas PC, Mattick JS. Extracellular DNA Required for Bacterial Biofilm Formation. Science. 2002;295(5559):1487-.

32. Okshevsky M, Regina VR, Meyer RL. Extracellular DNA as a target for biofilm control. Curr Opin Biotechnol. 2015;33:73–80.

33. Panlilio H, Rice CV. The role of extracellular DNA in the formation, architecture, stability, and treatment of bacterial biofilms. Biotechnol Bioeng. 2021;118(6):2129–41.

34. Goodman SD, Bakaletz LO. Bacterial Biofilms Utilize an Underlying Extracellular DNA Matrix Structure That Can Be Targeted for Biofilm Resolution. Microorganisms. 2022;10(2):466.

35. Lin Q, Sheng M, Tian Y, Li B, Kang Z, Yang Y, et al. Antibiofilm activity and synergistic effects of DNase I and lysostaphin against Staphylococcus aureus biofilms. Food Quality and Safety. 2024;8.

36. Tetz GV, Artemenko NK, Tetz VV. Effect of DNase and antibiotics on biofilm characteristics. Antimicrob Agents Chemother. 2009;53(3):1204–9.

37. Flemming H-C, Wingender J. The biofilm matrix. Nature Reviews Microbiology. 2010;8(9):623–33.

38. Hall CW, Mah TF. Molecular mechanisms of biofilm-based antibiotic resistance and tolerance in pathogenic bacteria. FEMS Microbiol Rev. 2017;41(3):276–301.

39. Harding CM, Hennon SW, Feldman MF. Uncovering the mechanisms of Acinetobacter baumannii virulence. Nat Rev Microbiol. 2018;16(2):91–102.

40. Sharples GJ, Ingleston SM, Lloyd RG. Holliday junction processing in bacteria: insights from the evolutionary conservation of RuvABC, RecG, and RusA. J Bacteriol. 1999;181(18):5543–50.

41. Bennett RJ, Dunderdale HJ, West SC. Resolution of Holliday junctions by RuvC resolvase: cleavage specificity and DNA distortion. Cell. 1993;74(6):1021–31.

42. Yamada K, Miyata T, Tsuchiya D, Oyama T, Fujiwara Y, Ohnishi T, et al. Crystal structure of the RuvA-RuvB complex: a structural basis for the Holliday junction migrating motor machinery. Mol Cell. 2002;10(3):671–81.

43. Okshevsky M, Meyer RL. The role of extracellular DNA in the establishment, maintenance and perpetuation of bacterial biofilms. Crit Rev Microbiol. 2015;41(3):341–52.

44. Bjarnsholt T, Kirketerp-Møller K, Jensen P, Madsen KG, Phipps R, Krogfelt K, et al. Why chronic wounds will not heal: a novel hypothesis. Wound Repair Regen. 2008;16(1):2–10.

45. Lebeaux D, Ghigo JM, Beloin C. Biofilm-related infections: bridging the gap between clinical management and fundamental aspects of recalcitrance toward antibiotics. Microbiol Mol Biol Rev. 2014;78(3):510–43.

46. Høiby N, Bjarnsholt T, Moser C, Bassi GL, Coenye T, Donelli G, et al. ESCMID guideline for the diagnosis and treatment of biofilm infections 2014. Clin Microbiol Infect. 2015;21 Suppl 1:S1–25.

47. Jakubovics NS, Shields RC, Rajarajan N, Burgess JG. Life after death: the critical role of extracellular DNA in microbial biofilms. Lett Appl Microbiol. 2013;57(6):467–75.

48. Koo H, Allan RN, Howlin RP, Stoodley P, Hall-Stoodley L. Targeting microbial biofilms: current and prospective therapeutic strategies. Nat Rev Microbiol. 2017;15(12):740–55.

